# Chimera-competent eXtra-Embryonic eNdoderm (XEN) cells established from pig embryos

**DOI:** 10.1101/2020.01.02.892802

**Authors:** Chi Park, Young Jeoung, Jun Uh, Kieun Park, Jessica Bridge, Anne Powell, Jie Li, Laramie Pence, Tianbin Liu, Hai-Xi Sun, Ying Gu, Yue Shen, Jun Wu, Juan-Carlos Izpisua Belmonte, Bhanu P. Telugu

## Abstract

In this article, we report for the first time the derivation and characterization of extra-embryonic endoderm (XEN) cells from primitive endoderm (PrE) of porcine (p) embryos. The pXEN cells can be reliably and reproducibly generated from parthenote, in vitro and in vivo derived embryos. The pXEN cells retained all the hallmarks of PrE including expression of canonical PrE and XEN cell markers (*GATA4, GATA6, SOX17, SALL4, FOXA2*, and *HNF4A*). Transcriptome analysis further confirmed their XEN cell origin. The pXEN cells when introduced into blastocyst stage embryo contributed to wide-spread chimerism including visceral yolk sac, chorion, as well as embryonic gut and liver primordium in the fetus. The pXEN cells were shown to be an efficient nuclear donor for generating cloned offspring. Taken together, pXEN cells fulfil a longstanding need for a stable, chimera-competent, and nuclear transfer-compatible porcine embryonic cells with applications for agriculture and medicine.

**Significance Statement:** We report for the first time, the derivation and characterization of extraembryonic endoderm (XEN) stem cells from porcine (p) embryos. The pXEN cells can be reliably and reproducibly derived from primitive endoderm precursors. When injected into blastocyst-stage embryos, the pXEN cells have contributed to wide-spread chimerism including visceral yolk sac, chorion of the extraembryonic membranes, as well as definitive endoderm of the fetus, primarily the embryonic gut and liver primordium. Additionally, these XEN cells have proven to be an efficient nuclear donor for generating cloned offspring. These newly discovered stem cells provide a novel model for studying lineage segregation, as well as a source for interspecies chimeras for generating endodermal organs, and for genome editing in livestock.

## Introduction

In mammals, delamination of primitive endoderm (PrE) from the inner cell mass (ICM) in the late blastocyst-stage embryo marks the second fate specification event (the first being the separation of trophectoderm (TE) from the ICM). The PrE differentiate into visceral endoderm (VE) and parietal endoderm (PE) that line ICM and TE, respectively(1). Together, the VE and PE generate yolk sac, the first placental membrane. The yolk sac serves as the main placenta in rodents until mid-gestation (d11.5), and performs several important functions including providing nutritional support, gas exchange, hematopoiesis, and patterning cues to the developing embryo. However, in non-rodent species including pig and humans, the yolk sac is short-lived (2, 3). Regardless, in all species the PrE does not contribute to the embryonic endoderm, which emerges later following gastrulation (4).

In culture, three types of stem cells can be established from the mouse embryo: embryonic stem cells (ESC) from the EPI, trophoblast stem cells (TSC) from TE, and XEN cells from PrE, which contribute to embryo proper, the placenta, and the yolk sac, respectively (5). The XEN cells can also be induced from ESC by overexpression of PrE-specific genes, *Gata-4, 6*(6, 7), or *Sox17*(8), or by treatment with growth factors(9). More recently, naïve extraembryonic endodermal cells (nEnd) resembling the blastocyst-stage PrE-precursors have been developed from the authentic mouse ESC (10). In rat, XEN cells established from blastocysts have different culture requirements and gene expression profiles compared to mouse XEN cells (11, 12). While, mouse XEN cells mainly contribute to the PE (13) in chimeras, the rat XEN cells contribute to the VE(12). It is unclear whether XEN cells from non-rodent animals (human and pig) have potency similar to mouse or rat (14). In this regard, the pig model can prove to be uniquely valuable in bridging the translational gap between rodents and humans.

Authentic ESC from pigs (p) have yet to be generated even after three decades of extensive investigation using conventional stem cell derivation conditions. The major reason for difficulties in the derivation of pESC is the instability of the pluripotent state(15, 16). Even though derivation of pESC from EPI cells has proven to be difficult, extraembryonic (ExE) cells within the early blastocyst outgrowths grow rapidly and outnumber the EPI cells, which can often be misinterpreted as epiblast cells(17). There are several reports describing pig EPI-like cells with properties similar to human ESC(18, 19). However, these observations are purely conjectural, only fulfilling minimal criteria of pluripotency, and lacking the deterministic *in vivo* demonstration of pluripotency(18, 20). Besides ESC, attempts to establish TSC and XEN cells from pig or other domestic animals has received little attention, and efforts to explore their potential is non-existent.

Here we describe the establishment of XEN cells from the PrE of the pig blastocysts. To-date these pXEN cells represent the only well characterized blastocyst-derived stem cell lines that can be readily and reproducibly established under conventional stem cell culture conditions. The pXEN cells are stable in culture, undergo self-renewal for extended periods of time, and contribute predominantly to yolk sac and at a minor level to embryonic endoderm (gut) in chimeras, and can serve as nuclear donors to generate live offspring.

## Results

### *In vitro* derivation and expansion of primary PrE outgrowths

A central assumption behind the failure to establish pESC is a rapid loss of pluripotency in primary outgrowths(15). The whole blastocyst explants following attachment became flattened and spread out within 2 days of culture (Fig. 1a). As primary outgrowth expanded, TE cells began to first emerge and then underwent dramatic morphological changes, becoming larger and flatter, and soon-after undergoing apoptosis (Fig. 1a). After 5 days, a population of round and dispersed epithelial cells emerged as a discrete cell layer bordering the ICM (hereafter called “EPI”) cells (Fig. 1a). Majority of EPI cells were SOX2 positive (18/21) but only a few co-expressed NANOG (4/21) (Fig. 1b), similar to the staining pattern observed in the blastocyst (Fig.1c). Notably, the large round cells, initially considered as TE cells stained positive for GATA6 (9/12) and CK18 but lacked CDX2 expression (Fig. 1b). The expression of GATA4, a later marker of the PrE-was also detected in few small round cells (4/7) (*SI Appendix, Fig. S1a*), confirming two distinct PrE progenitors expressing GATA factors in primary outgrowths. These subpopulations, small and large PrE were distinguishable based on cell morphology and by their expression of CK18 (*SI Appendix, Fig.S1b*). Although initial explants could be established from early blastocysts (day 5-6), late blastocysts (fully expanded or hatched, day 7-8), where the ICM and TE lineages were discernable (Fig. 1c; *SI Appendix, Fig. S1c*) established stable PrE populations (Fig. 1d-e) and were used for further studies.

**Figure 1.**
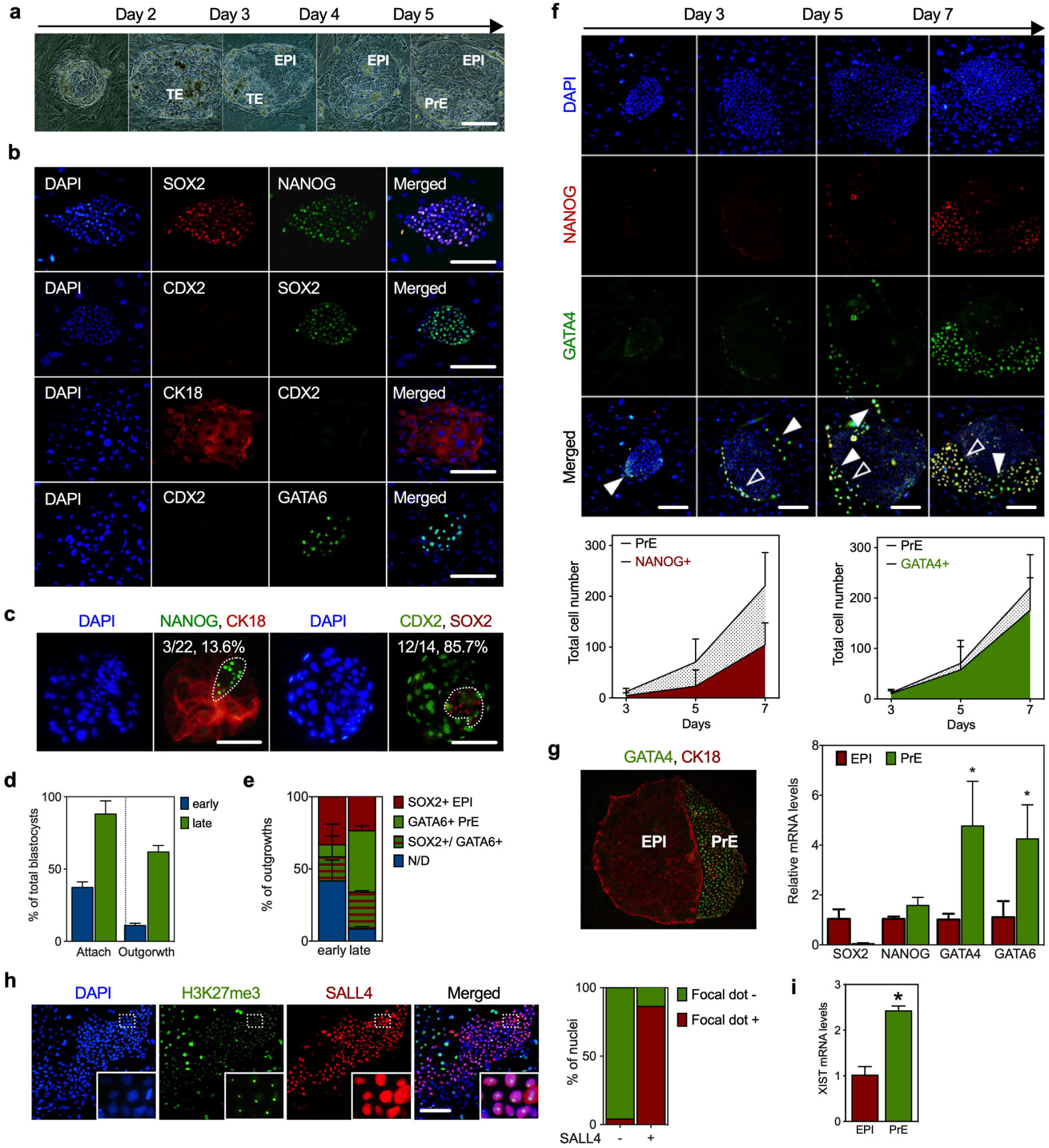
Distinct subpopulations arise from the porcine blastocyst outgrowths. (a) Phase contrast images depicting morphologies of embryonic outgrowths from days 2 to 5 in culture. In the figure EPI, TE and PrE stands for epiblast, trophectoderm and primitive endoderm, respectively. (b) Immunostaining for key transcription factors, SOX2 and NANOG (ICM), CDX2 and CK18 (TE), and GATA6 (PrE) in the primary outgrowth at day 3 after explants. (c) Representative immunofluorescence images of late blastocyst (ICM in dotted circle). In the figure, fraction of cells and percentage of cells that stained positive for NANOG or SOX2 was shown. (d) The bar graph showing the attachment and outgrowth rates of early and late blastocysts. (e) Frequencies of SOX2- and GATA6-positive cells in outgrowths. N/D: not detected. (f) Representative immunostaining (top) and quantitation (bottom) of the number of NANOG or GATA4 positive nuclei in primary outgrowths cultured for 7 days. Open and solid arrows indicate NANOG/GATA4 co- positive and GATA4 positive only cells, respectively. (g) Representative fluorescence images of CK18 and GATA4 of a Day 7 primary outgrowth (right). Comparison of the transcriptional levels of selected lineage marker genes between PrE cells and EPI cells by qPCR; *, p<0.05 according to unpaired *t* test; error bars represent ± SEM (n=3) (left). ACTB was used as an endogenous control. (h) The expression of H3K27me3 and SALL4 in day 7 primary outgrowth (right). Inset shows the zoom-in of the dashed box. The bar graph showing the quantitation of the percentage of H3K27me3 focal dots in SALL4 positive or negative cells (left). In all images, nuclei were counterstained with DAPI (blue). Scale bar: 100µm. (i) The relative XIST mRNA levels in PrE cells compared to EPI cells; *, p<0.05 according to unpaired *t* test; error bars represent ± SEM (n=3). ACTB was used as a loading control.

Initially, NANOG or GATA4 positive (+) cells were mostly undetectable, but cytoplasmic GATA4 expression appeared in the periphery of the early ICM outgrowths by d3 of culture (Fig. 1f). Intriguingly, NANOG/GATA4 co-positive cells that lined the side of EPI outgrowth gradually increased by 5 days, and by d 7, > 90% of GATA4+ cells co-expressed NANOG (Fig. 1f). In contrast, the expression of NANOG was detected in few, if not at all in EPI cells, while the SOX2 expression was progressively decreased with time, indicating the loss of pluripotency (*SI Appendix, Fig. S1D*; Fig.1e,1g). Besides GATA factors, SALL4(21) a key stemness marker of XEN cells was expressed in the nuclei of the PrE cells that had a small and compacted appearance. A large fraction (∼75%) of SALL4+ cells had nuclear foci of intense histone 3 lysine 27 trimethylation (H3K27me3), a hallmark of the inactive X in female outgrowth(22) (Fig. 1h). Consistent with this observation, *XIST* levels were 2-fold higher in SALL4+ PrE cells than EPI cells (Fig. 1i), which reflects the lineage specific dynamics of H3K27me3 accumulation on the X-chromosome, and could be the consequence of the co-expression of *SALL4*(21).

### Cellular properties and molecular signature of pig XEN cells

Self-renewal of XEN cells is dependent on *Sall4* expression(21). The emergence of distinct SALL4+ PrE population in primary outgrowths have prompted us to attempt derivation of pXEN cells. After 7-9 days of culture, PrE cells began to emerge in primary outgrowths and could be clearly demarcated based on their morphology and allowing for easy dissociation from the EPI cells (*SI Appendix, Fig.S2a*). Both EPI and PrE colonies displayed a distinct morphology following serial passages (Fig. 2a). Consistent with previous findings, the EPI colonies underwent spontaneous differentiation toward a fibroblast- or neuron-like appearance by passages 5-7. The colonies from PrE-derivatives on the other hand, were more stable in culture. The colonies were propagated as flattened colonies and passaged as clumps by mechanical or enzymatic dissociation (Fig. 2b), but did not survive passage as single cells, even when treated with ROCK inhibitor Y-27632 (Fig. 2b; *SI Appendix, Fig.S2b*). Following sub-passage, the PrE colonies initially appeared as a homogenous colony of cells and grew as a single sheet monolayer. Upon serial passaging two distinct populations emerged; a cobble-stone morphology in the center of colony, and an epithelial sheet-type cells at the borders of the colony (*SI Appendix, Fig.S2c*). The cells at the periphery were strongly alkaline phosphatase (ALP) positive (Fig. 2c) and exhibited rapid proliferation as confirmed by PCNA staining (*SI Appendix, Fig.S2c*). The density of the feeder cells influenced the colony stability with the optimal densities ranging from 3-4 × 10^4^ cells per cm^2^. Lower feeder densities (< 2 × 10^4^ cells/cm^2^) resulted in differentiation of cells with the expression of VIMENTIN (Fig. 2d), and at high density (>1 × 10^5^ cells/cm^2^), the cultures were more closely packed and showed reduced replating efficiency. The cells expressed PrE-specific markers (GATA4, GATA6, SOX17, SALL4, FOXA2, and HNF4A) with no expression of pluripotent markers (OCT4, SOX2, and NANOG) (Fig. 2e; *SI Appendix, Fig.S2e*). Notably, NANOG was no longer detected upon passaging indicating a possible role for NANOG only in early PrE specification. While CDX2 is not detectable, other TE-markers EOMES and GATA3 were expressed, consistent with the role of the latter in endodermal specification. Taken together, the molecular signature confirmed the established colonies as XEN cells.

**Figure 2.**
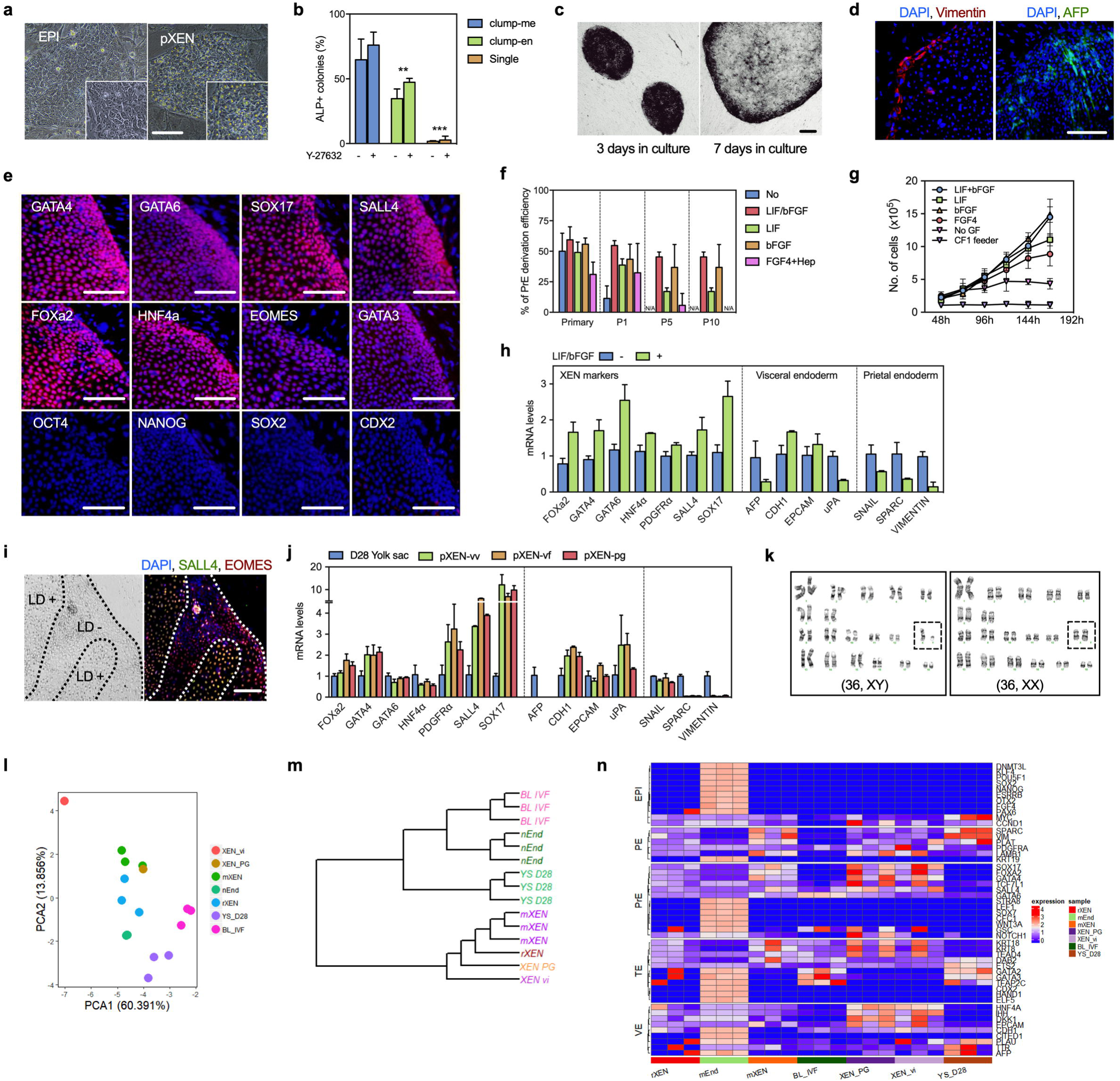
Establishment and characterization of pXEN cells. (a) Representative bright-field images of EPI- derived primary colonies, and PrE-derived XEN cells at passages 3-5. (b) Efficiency of colony formation of pXEN cells passaged as clumps or single cells. The colony forming activity were greatly impaired when dissociated as single cells. Cells were passaged as clumps by mechanical (clumps-me) or enzymatic dissociation (clumps-en) with Accutase. (c) Alkaline phosphatase (ALP) staining of an *in vivo*-derived pXEN cells (Xv#9) after culturing for 3 and 7 days. (d) Representative fluorescence images of VIMENTIN (red) and AFP (green) (e) Expression of the indicated markers in pXEN at passages 30-35. (f) Effect of growth factors supplementation on PrE derivation. pXEN cells were seeded onto a 6-well-plate seeded containing a density of 5□×□10^4^ feeder cells per cm^2^, and (g) cell number estimated 48 h following passage. Data are is presented as means□± s.d. (n = 3). (h) qPCR analyses of total RNA isolated from pXEN cells grown in either the presence or absence of LIF/bFGF for 4 days. ACTB was used as a loading control. The values are represented as mean□±□s.d. (n = 3). (i) Representative images of pXEN cells show the expression of stem cell marker, SALL4 (green) that are significantly reduced in the cells that had lost lipid droplet. Scale bar: 100µm. (j) qPCR analysis of pXEN cells derived from different embryonic origins. ACTB was used as a loading control. The values are represented as mean□±□s.d. (n = 3). (k) Representative karyotypic analysis of pXEN cell lines, with numbered chromosomes. (l) RNA-seq analysis of pXEN cells and comparison with analogous derivatives. Data from pig XEN cell lines as well as published data on related cell lines (mouse and rat XEN cells) were included in the comparison. Principal component analysis (PCA) plot of two pXEN cells and other samples. Upper inset shows the color code for each cell type, lower inset shows a separate PCA of only pig vs. mouse vs. rat XEN cells. (m) hierarchical clustering of pXEN and related samples. (n)Heatmap comparison of selected XEN-associated extraembryonic endodermal (ExEn) marker gene expression of all samples.

We tested the growth factor requirements of pXEN cells based on observations from mouse(23). Withdrawal of either LIF, bFGF or both, had no impact on primary PrE induction. However, in the omission of both, the cells failed to expand into stable cell lines confirming the growth factor responsiveness (Fig. 2f). The colonies that arose in the LIF or FGF4 alone did not proliferate as rapidly as cells cultured with either bFGF, or both LIF and bFGF (Fig. 2g). Omission of both growth factors resulted in a dramatic reduction in colony formation, with low expression of XEN marker genes *FOXa2, GATA4, GATA6, HNF4a, PDGFRa, SALL4* and *SOX17*, and high expression of VE- (*AFP* and *UPA*), and PE-genes, (*SNAIL, SPARC*, and *VIMENTIN*), consistent with spontaneous differentiation (Fig. 2h). The XEN cells can be stably maintained in serum-free N2B27-based defined medium with lower degree of cellular differentiation and expression of VE- and PE-related genes, however requiring a longer cell doubling time (*SI Appendix, Fig.S2f*; 2g). One interesting finding is the presence of characteristic lipid droplets in the cytoplasm of pXEN cells (Fig. 2a), which readily disappeared when plated in the absence of growth factors or feeder cells with a concomitant loss of SALL4 expression, but no change in EOMES expression (Fig. 2i). Although little is known about the mechanisms mediating the presence of lipid droplets, this feature could be leveraged as a non-invasive marker of SALL4+ cells.

Based on these preliminary trials, we established putative XEN cell lines from *in vivo*-developed (vi, n=4), *in vitro*-fertilized (vf, n=13), and parthenogenetically activated (pg, n=14) porcine blastocysts that exhibited stable morphology and marker expression, irrespective of the origin of embryos (Fig. 2j). The pXEN cells were maintained with proliferative potential in culture for extended passages (>50 passages), and were karyotypically normal (Fig. 2k). Transcriptomic analysis of pXEN cells expressed characteristic XEN cell repertoire and clustered closely with rodent XEN cells (Fig. 2l, 2m, 2n). Importantly, no teratoma development was observed in any recipient mice transplanted with the six robust pXEN cell lines ranging from 1 × 10^6^ to 10^7^ cells (*SI Appendix, Table S2*) indicating that all injected pXEN cells were committed and not pluripotent cells.

### Contribution of pXEN Cells to Chimeras

Mouse XEN cells contribute to PE, whereas rat XEN cells incorporate into both VE and PE lineages in chimeras(12, 23). Given these disparities, we evaluated the properties of pXEN cells in chimera studies (Fig. 3a). To facilitate lineage tracing, we generated a novel reporter pXEN cell line by knocking-in a constitutive human UBC promoter driven GFP reporter downstream of the pCOL1A1 locus (hereafter, pCOL1A:GFP) using CRISPR/Cas system as previously described(24) (*SI Appendix, Fig.S3a*). Labeled pXEN (Xnt ^pCOL1A:GFP^ #3-2) cells were injected as single cells or 5-10 cell clumps into parthenogenetic embryos at the morula (Day 4) or early blastocyst stages (Day 5). Cells injected as clumps efficiently integrated into host embryos (77.3 to 85.7%) than individual cells (37.5 to 47.4%); and cells injected at the blastocyst stage showed better incorporation into ICM (85.7%) than injection at morula stage (77.3%) (*SI Appendix*, Table S3). To evaluate *in vivo* chimeric development, pXEN cells were similarly injected as clumps into host blastocysts (n=109) and the resulting re-expanded blastocysts following overnight culture (n=94) were transferred into 3 recipient sows (Fig. 3b). A total of 25 fetuses (27%) were retrieved from 2 recipients on days 21(Fig. 2b). Among the recovered fetuses, the injected GFP+ cells were found in the yolk sac (6/9) and the fetal membranes (5/9), and a small group of GFP+ cells were observed in one embryo (1/9) (Fig. 3b). Notably, GPF+ cells extensively contributed to yolk sac in two chimeras (XeC#2-3 and XeC#2-4) with a moderate signal in allantochorion (Fig. 3c). The GFP+ cells observed in embryos were from pXEN cells and not due to auto-fluorescence as confirmed by genomic PCR. Quantification of GFP+ cells by qPCR confirmed XEN cell chimerism at 1.7% in 2 embryos, and at 12.9% in the yolk sac, and 8% in the allantochorion, signifying active integration and proliferation of pXEN cells during embryogenesis (Fig. 3d). As shown in Fig. 3e, immunostaining with the anti-GFP antibody identified GFP+ cells in the embryonic gut of 3 chimeric fetuses (XeC#1-2, XeC #2-3, and XeC #2-6). The GFP+ donor cell population integrated predominantly into the visceral endodermal layers, but rarely into the outer mesothelial layers or endothelial cells in the yolk sac (Fig. 3e middle; *SI Appendix, Fig.S*3c), and to a minor extent populated amnion, allantois, chorion (Fig. 3e; *SI Appendix, Fig.S*3d), and gut endoderm (*SI Appendix*, Table S5). Overall, the chimerism frequency of the pXEN cells was rather high (60%).

**Figure 3.**
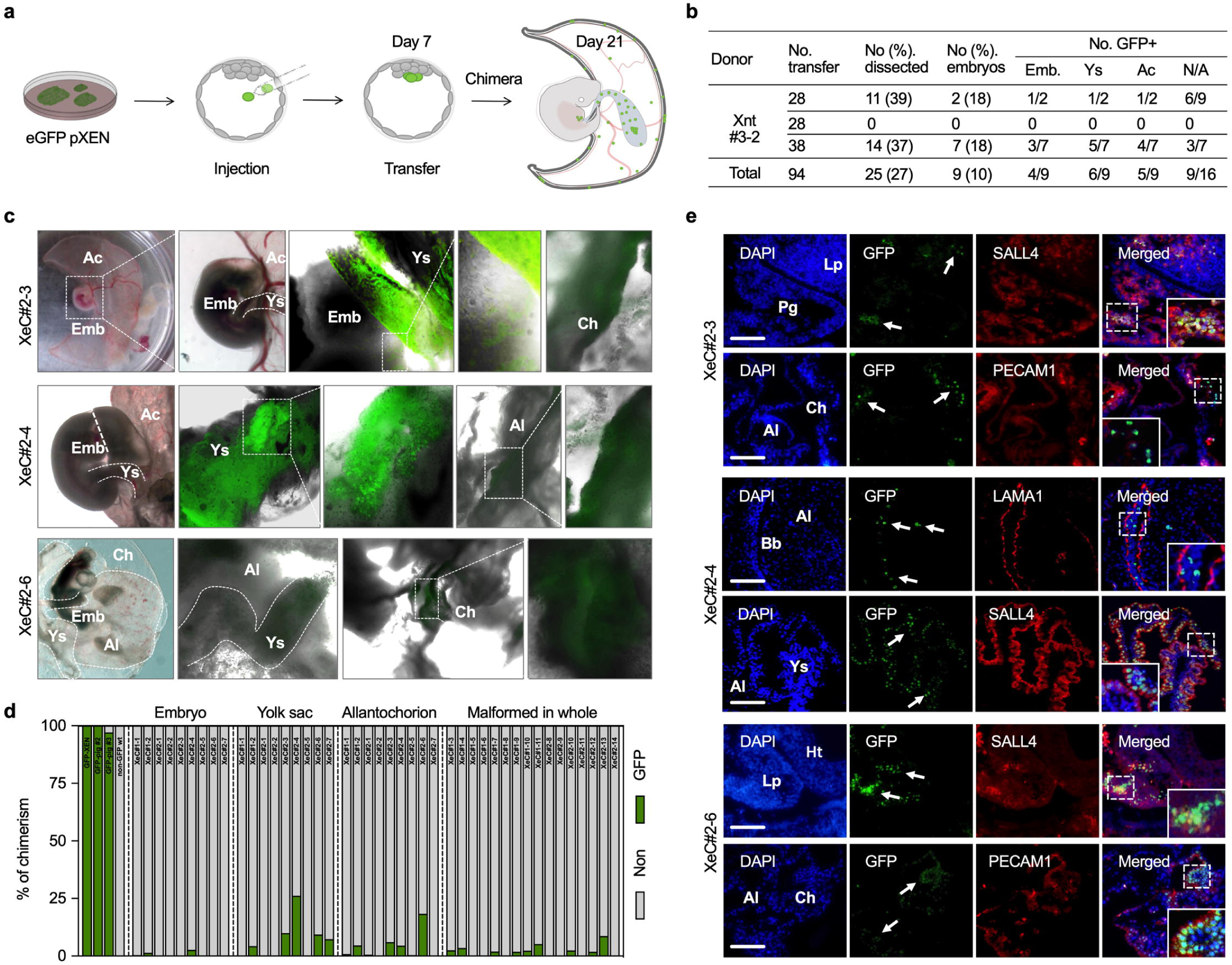
Chimeric contribution of pXEN cells to embryonic and extraembryonic lineages in post-implantation Day 21 embryos. (a) Schematic representation of the chimera assay. (b) Table presents a summary of chimera experiments performed by injection of pXEN cells into blastocysts. In the Table, Ys: yolk sac; ExE: extraembryonic membranes; N/D: not defined (severely retarded fetuses with no fetal or yolk sac parts); and “*” stands for the embryos at the pre-attachment stages (spherical or ovoid). (c) Representative bright field and fluorescence merged images of normal (XeC#2-3 and XeC#2-4) and retarded (XeC#2-6) fetuses at day 21 of gestation. Yolk sac outlined by the dashed line, and enlarged view of the region marked by the dashed box is shown in the right. In the figure Al stands for allantois; Ch, chorion; Emb, embryo; Ys, yolk sac. (d) Bar graph representing percent contributions of GFP-XEN in chimeras determined by qPCR; *, p<0.05 according to unpaired *t* test; error bars represent ± SEM (n=3). (e) Representative sagittal or transverse sections of fetuses showing dual immunofluorescence staining for GFP (green) and SALL4 or PECAM1 (red) in embryos; the arrows indicate GFP-positive cells derived from injected pXEN cells in sections. Inset are zoom-in magnified images of the dashed box. Nuclei were stained with DAPI (blue). Al, allantois; Ch, chorion; Emb, embryo; Lp, liver primordium; Pg primitive gut; Ys, yolk sac; Am, amnion; Hp, heart primordium; So, somite. Scale bar: 100µm.

### Generation of viable cloned offspring from pXEN cells via SCNT

In an effort to test the utility of pXEN cells as nuclear donors, we performed SCNT with the pXEN cells used in the chimera assay (above), alongside previously published crossbred knock-out fetal fibroblasts (FF ^NGN3-/-^) as controls(25). A total of 222 cloned embryos reconstituted from pXEN (Xnt ^pCOL1A:GFP^ #3-2, n=61) and FF (n=161) were co-transferred into two surrogate gilts to exclude confounding variables associated with recipient animals affecting the outcome. Following embryo transfers, one pregnancy was established, and delivered 8 cloned piglets at term. Three of the 8 piglets were GFP positive and black coated (4.9%) confirming the COL1A:GFP Ossabaw XEN cell origin, while 5 piglets were white coated and GFP negative from the control fibroblasts (3.1%) (Fig. 4a). As expected, the piglets exhibited ubiquitous expression of GFP in all tissues (Fig. 4b). The genotype of the offspring was confirmed by PCR (Fig. 4c). In addition to this, we performed multiple rounds of SCNT with FF ^pCOL1A:GFP^ (#3) from which the XEN cells were derived. Despite being genetically identical, no offspring were obtained from founder GFP fibroblasts, but the derived XEN cells served as efficient donors in SCNT.

**Figure 4.**
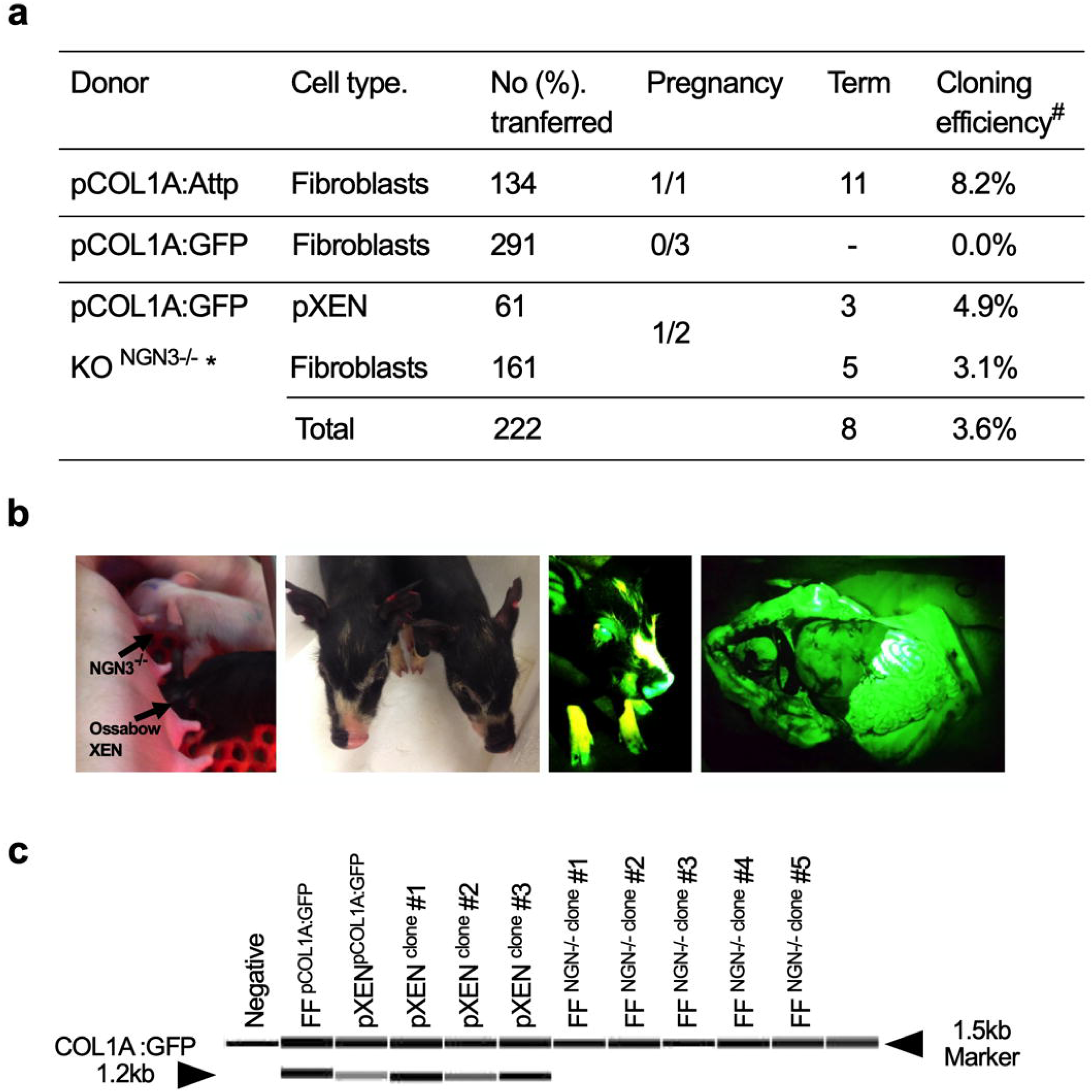
Generation of viable cloned piglets using pXEN or fibroblasts. (a) Summary of SCNT experiments. ^#^Cloning efficiency was obtained by calculating total no. fetuses or piglets / total no. embryos transferred. ^$^data obtained from our previous study. *NGN3^-/-^ cells originated from our previous report^25^. All the fetal fibroblasts and pXEN cells with the exception of NGN3^-/-^ cells used as SCNT donors were derived from the same fetus (female Ossabow fetal fibroblast #6). (b) Representative images showing 10 days old NGN3 KO white (outbred)- and XEN Black (Ossabaw)-coated littermates. The fluorescence images of live GFP+ piglets and whole organs taken with blue light illumination showing ubiquitous expression of GFP transgene, and confirming the pXEN cell as nuclear donors. (c) A representative digital gel image of the 1.2-kb amplicon with primers within and outside of the targeting vector confirming site-specific knockin was generated by Fragment Analyzer.

## Discussion

Establishment of pESC from embryonic explants without major chemical interventions has largely been unsuccessful despite nearly three decades of investigation. As shown by multiple groups, the EPI fraction of the primary explants fail to proliferate, and the cultures are rapidly overtaken by proliferating ExE cells. That said, there were no published reports that temporally followed the fate of the ExE derived lines in culture, nor have they been adequately characterized; although, the equivalent lines from mouse have been thoroughly characterized. This report for the first time takes a systematic and in-depth look at the derivation, establishment, and characterization of XEN cells from PrE.

During early mouse embryo development, NANOG is expressed in EPI cells and excluded from GATA4+ PrE cells in embryo(26, 27). This seems counterintuitive given the mutual antagonism between NANOG and GATA4 that facilitate key cell-fate decisions between EPI and PrE, respectively(28). Indeed, several lines of evidence support the expression of NANOG in pig hypoblast(29, 30), which is contrary to the mouse model. Emergence of PrE population with co-expression of GATA4/NANOG appears to represent an early step in PrE specification, highlighting mechanistic differences in early lineage specification between mouse and pig. That said, the establishment of pXEN cells, culture characteristics, and the resulting molecular signatures (including high expression of *FOXa2, GATA4, GATA6, HNF4a, PDGFRa, SALL4* and *SOX17*) are shared with rodent models, with the exception of failure to establish XEN cells in FGF4-based medium, and intolerance to dispersal as single cells.

Generation of embryonic chimeras has been considered as the most stringent test of stem cell differentiation potential *in vivo(31)*. This study demonstrates that despite the lack of pESC, it is possible to generate embryo-derived stem cell lines with PrE-like properties as confirmed by lineage-restricted plasticity in the resulting chimeras, which were not irrevocably fixed (e.g., yolk sac, placenta, gut endoderm)(4). This indicates that the pXEN cells are in a less committed endodermal naïve state. In the pig, freshly isolated ICMs are capable of widespread tissue contribution, including germline colonization in chimeras(32). Despite this, the pluripotent EPI or iPS cells were preferentially engrafted into extraembryonic tissues(33-35). It is likely that in the absence of defined conditions, embryonic outgrowths are unstable and transition to a XEN-like state(36). Future chimera trials will be performed in embryos that lack key gate-keeper genes (for e.g., SALL4), where the relative contribution of pXEN cells to embryonic and ExE endodermal lineages are expected to be higher when compared to current experiments performed with wild type embryos.

*In vivo* generation of human organs via interspecies chimeras between human and pigs via blastocyst complementation has been acknowledged as a source of donor organs for life-saving regenerative medicine applications(37-39). Evidence gathered in the present study demonstrates the engraftment potential of pXEN cells with lineage restricted cell fate. When such experiments are performed with human XEN cells, the potential contribution to endodermal organs will provide an on-demand source of human endodermal cells in pig hosts. Our present findings make the use of pXEN cells a particularly attractive choice to generate tissue-specific chimeras for endodermal organs, while limiting unwanted contribution to undesirable organs (e.g., germ cell or neural lineage) in interspecies chimeras-a likely outcome with the use of ESC/iPS cells(40, 41).

Another advantage of the pXEN cells is the competency to generate live animals via SCNT. This is especially attractive in complex genome editing and genetic engineering applications where long-life span in culture is desirable. As evidenced from this study, genetically modified fibroblast cells failed to generate live offspring, whereas, the pXEN cells derived following cloning of the FFs were able to generate live offspring at a relatively high efficiency (4.9%). One potential explanation is the epigenetic disruption caused by transfection that may *have* compromised embryonic development. The pXEN cell derivation processes, has potentially reset the genome to a state that allows full-term development. It remains to be seen, if this could be applicable to other cells which failed to generate live offspring. Taken together, we argue that the derivation of pXEN cells fulfils a longstanding need in the livestock genetics for a stem cell line of embryonic origin that can be reliably and reproducibly generated, are stable in culture, have the potential to contribute to chimeras, and are a good source for creating cloned animals.

## Materials and Methods

More detailed information regarding the materials and methods are available in SI Appendix. All methods for derivation and characterization has been outlined in the Supplementary Appendix. A total of 12 RNA-seq data sets generated in this study have been deposited in the CNSA (https://db.cngb.org/cnsa/) of CNGBdb with accession code CNP0000388, and also NCBI Gene Expression Omnibus (GEO; http://www.ncbi.nlm.nih.gov/geo) under accession number GSE128149.

## Acknowledgments

This research was supported by AFRI Grants# 2015-67015-22845 and 2018-67015-27575 from the USDA National Institute of Food and Agriculture, and 1 R01 HD092304-01A1 from NICHD, NIH to BT; Guangdong Provincial Key Laboratory of Genome Read and Write (No. 2017B030301011), Guangdong Provincial Academician Workstation of BGI Synthetic Genomics (No. 2017B090904014), Shenzhen Engineering Laboratory for Innovative Molecular Diagnostics (DRC-SZ[2016]884) and Shenzhen Peacock Plan (No.. KQTD20150330171505310)

