## Supplementary Appendix for "Chimera-competent eXtra-Embryonic eNdoderm (XEN) cells established from pig embryos"

**The PDF file includes:**

- Materials and Methods
- Appendix Table s1. Primers and Antibodies
- Appendix Table s2. Teratoma assay for determining the potency of pig XEN cells
- Appendix Table s3. Integration of pXEN cells to pig blastocysts
- Appendix Table s4. A detailed analysis of distribution GFP+ cells in D21 pig chimeras
- Appendix Table s5. In vitro development of pig cloned embryos
- Appendix Figure s1. Self-renewal of extra-embryonic endoderm XEN cells.
- Appendix Figure s2. Distinct subpopulations arise from the blastocysts outgrowth.
- Appendix Figure s3. Chimeric contributions of pXEN cells in embryo.

### Materials and Methods

#### Animal Experimental Assurance

All experiments involving live animals were performed in accordance with the approved guidelines of the Beltsville ARS and Thomas D. Morris Inc., Institutional Animal Care and Use Committee (IACUC). All experimental protocols involving live animals were approved by the IACUC committee.

#### Establishment and maintenance of pig XEN cells

Embryonic explants and XEN cells were cultured on a feeder layer of early passage (n=3) CF-1 mouse embryonic fibroblasts (MEF) cells mitotically inactivated by treatment with mitomycin-C (3 hr, 10 µg/mL). A day before seeding the embryos or XEN cells, the feeders were plated in MEF medium based on high-glucose Dulbecco's modified Eagle medium (DMEM; Gibco) supplemented with 10% (v/v) fetal bovine serum (FBS; HyClone) on 0.1% (v/v) gelatin-coated four-well plates (Nunclon) at a density of 3-5 × $10^5$ cells per $cm^2$. At least 2 hr before the start of the experiment, the MEF medium was aspirated and replaced with 'standard ES medium' which included DMEM/ Nutrient Mixture Ham's F12 (DMEM/F-12; Gibco) supplemented with 15% ES-qualified fetal calf serum (FCS; HyClone), 1 mM sodium pyruvate, 2 mM L-glutamine, 100 units/mL penicillin-streptomycin, 0.1 mM 2-β-mercaptoethanol, 1% non-essential amino acids (NEAA; Gibco), with various combination of growth factors; 10 ng/mL human recombinant leukemia inhibitory factor (hrLIF; Milipore) and 10 ng/mL human recombinant basic fibroblast growth factor (hrbFGF; R&D Systems). Other media combinations that were tested include RPMI 1640 or N2B27 serum free medium (1:1 ratio of DMEM/F12 and Neurobasal medium plus N2 and B27, all from Gibco), with a combination of 5 ng/mL LIF and/or 10 ng/mL bFGF, or 25 ng/mL human recombinant fibroblast growth factor 4 (hrFGF4; R&D Systems) and 1 μg/mL heparin[1]. Following initial plating, attachment and outgrowth development, the medium was refreshed on d 3, followed by media exchange every 2 days. After 7-8 days of culture, the primary outgrowths were mechanically dissociated into small clumps, and transferred onto fresh feeders for passaging. The pXEN cells were cultured at 38.5°C in 5% $O_2$ and 5% $CO_2$, with the culture medium being refreshed every other day and passaged at 1:20 every 7-8 days. Cells were passaged as clumps by gentle pipetting following 10 min digestion with Accutase (Gibco). Before routine passaging and freezing, cells were cultured with Rho Kinase (ROCK) inhibitor Y-27632 (10 μM; StemCell Technologies) at least 2 hr prior to dissociation[2]. Each XEN cell line was frozen in FBS based medium supplemented with 8% (v/v) DMSO and recovered with high viability. In order to determine chromosomal stability in long term culture, cytogenetic analysis was performed by Cell Line Genetics.

**Alkaline phosphatase staining**

The cells were fixed with 4% (w/v) paraformaldehyde for 3 min at room temperature (RT) and were washed three times with DPBS. Alkaline phosphatase (ALP) staining was performed with a BCIP/NBT Alkaline Phosphatase Color Development Kit following the manufacturer's instructions. The cells were examined using an inverted microscope.

***In vitro* differentiation of XEN cells into parietal or visceral endoderm:**

The pXEN cells were differentiated by means of embryoid body (EB) formation and treatment with small molecules and factors as previously described[3]. pXEN cells were dissociated as clumps, washed, and resuspended in medium (DMEM/F12 plus 15%

FBS) as hanging drops on the lid of a 60 mm dish, and cultured for 5 days, during which time spheroids were formed. To direct pXEN cells differentiation into either visceral endoderm (VE) or parietal endoderm (PE), accutase-dissociated single cells ($2 \times 10^5$ cells per $cm^2$) were seeded onto a laminin- or fibronectin-coated 6 well plate in N2B27 medium supplemented with the respective differentiation factors and/or chemicals. For example, for differentiation into VE, the cells were treated with CHIR99021 (10 μM, STEMCELL Technologies Inc.) and BMP4 (50 ng/mL, R&D) for activating Wnt/β-catenin pathway; for differentiation into PE, Folskolin (50 μM) and dbcAMP (1mM) for activating the cyclic adenosine monophosphate (cAMP) signaling pathway were utilized. Differentiation medium was replaced every two days, and cells were processed for analysis on day 12.

**Methods for embryo production and manipulation**

The *in vivo* and *in vitro* embryo production were performed as described previously[4, 5]. For generating parthenote, *in vitro* fertilized embryos, and for performing somatic cell nuclear transfer (SCNT), cumulus-oocyte complexes were purchased from a commercial supplier (DeSoto Biosciences). After *in vitro* maturation, the cumulus cells were removed from the oocytes by gentle pipetting in a 0.1% (w/v) hyaluronidase solution. Briefly, for *In vitro* fertilization (IVF), pre-diluted fresh semen (Duroc; Progenes) was centrifuged twice at 200 g for 3 min in DPBS containing 0.2% BSA. The sperm pellet was adjusted to a concentration of $2 \times 10^5$ sperm per mL and co-incubated with matured oocytes in modified Tris-buffered medium containing 0.4% BSA for 5 hr in a humidified atmosphere (5% $CO_2$ in air). Following three washes, putative zygotes were cultured and maintained in PZM3 medium in a low oxygen air (5% $O_2$ and 5% $CO_2$ in air). For obtaining *in vivo* embryos, donor animals were synchronized using Regumate and artificially inseminated at 12 and 24 hr following the observation of first standing estrus. On days 5-7 post-insemination, *in vivo* embryos were recovered by flushing oviduct with 35 ml of TL-Hepes buffer containing 2% BSA under general anesthesia. For SCNT, fetal fibroblasts (FF) were synchronized to the G1/G0-phase by serum deprivation (DMEM with 0.2% FCS) for 96 hr, and pXEN cells were mitotically arrested by serum free medium (N2B27 with 1% BSA) for 48 hr followed by incubation with aphidicolin (0.1 μM) for 12 hr. Enucleation was performed by aspirating the polar body and the MII metaphase plates using a micropipette (Humagen, Charlottesville, VA, USA) in 0.1% DPBS supplemented with 5 μg/mL of cytochalasin B. After enucleation, donor cells were placed into the perivitelline space of an enucleated oocyte. Fusion of cell–oocyte couplets was induced by applying two direct current (DC) pulses (1-sec interval) of 2.1 kV/cm for 30 μs using a ECM 2001 Electroporation System (BTX). After fusion, the reconstituted oocytes were activated by a DC pulse of 1.2 kV/cm for 60 μs, followed by post-activation in 2 mM 6-dimethylaminopurine for 3 hr. After overnight culture in PZM3 with a histone deacetylase inhibitor Scriptaid (0.5 μM), the cloned embryos were surgically transferred into the oviduct. Parthenogenetic embryos were produced by the activation procedures used for SCNT.

**Embryo Transfer**

The surrogate recipients were synchronized by oral administration of progesterone analog Regumate for 14-16 days. Animals in natural estrus on the day of surgery were used as recipients for SCNT embryo transfers (into oviduct), and at days 5-6 after natural heat were used for blastocyst transfer (into uterus) for generating chimeras.

Surgical procedure was performed under a 5% isofluorane general anesthesia following induction with TKX (Telazol 100 mg/kg, ketamine 50 mg/kg, and xylazine 50 mg/kg body weight) administered intramuscularly. Pregnancies were confirmed by ultrasound on day 27 following transfer. Cloned piglets were delivered at day 117 of pregnancy by natural parturition.

**RNA and DNA preparations**

For isolation of genomic DNA (gDNA) from cells and tissues, the QIAamp mini DNA Kit (Qiagen) was used according to the manufacturers' instructions. Total RNA was isolated using Trizol plus RNeasy mini kit (Qiagen) and mRNA from individual blastocysts was extracted using the Dynabeads mRNA Direct Kit (Dynal Asa). Synthesis of cDNA was performed using a High Capacity cDNA Reverse transcription kit (Applied Biosystems according to the manufacturers' instructions. The QIAseq FX Single Cell RNA Library kit (Qiagen) was used for Illumina library preparation and transcriptomics analysis.

**qPCR:** Relative quantification of mRNA levels was carried out using SYBR Green technology on an ABI 7500 Fast Real-Time PCR system (Applied Biosystems). The thermal-cycling conditions are: 20 s at 95°C followed by 40 cycles of 3 s at 95°C and 30 s at 60°C. The primers were designed to yield a single product without primer dimerization. The amplification curves for the selected genes were parallel. All reactions were performed from three independent biological and two technical replicates. Two reference genes, ACTB and YWHAG were used to normalize all samples and the relative expression ratios were calculated via the $2^{-\Delta\Delta Ct}$ method[6]. The primers used in qPCR are listed in sTable 1.

**Data access**

A total of 12 RNA-seq data sets generated in this study have been deposited in the CNSA (https://db.cngb.org/cnsa/) of CNGBdb with accession code CNP0000388, and also NCBI Gene Expression Omnibus (GEO; http://www.ncbi.nlm.nih.gov/geo) under accession number GSE128149.

**Transcriptomics Analysis**

RNA-seq reads were mapped to the pig reference genome (Sscrofa11.1) using HISAT2[7] (version 2.0.4) with parameters "hisat2 --sensitive --no-discordant --no-mixed -I 1 -X 1000" and to the reference cDNA sequence using Bowtie2[8] with parameters "bowtie2 -q --sensitive --dpad0 --gbar 99999999 --mp 1,1 --np 1 --score-min L,0,-0.1 -I 1 -X 1000 --no-mixed --no-discordant-p 1 -k 200". Then the expression levels of each gene were calculated by the fragments per kilobase of exons per million fragments mapped (FPKM) using RSEM[9] with parameters "rsemcalculate-expression --paired-end -p 8" based on the result of Bowtie2. The data of mouse and rat XEN cells were downloaded from GSE106158[10] (mouse: GSM2830587, GSM2830588 and GSM2830589; rat: GSM2830591, GSM2830592 and GSM2830593) and the gene expression levels were calculated in the same way (the mouse and rat reference genome used were GRCm38.p6 and Rnor_6.0, respectively). The expression levels of mouse nEnd were downloaded from GSE10742[11] (GSM271163, GSM271164 and GSM271165). Then the expression levels of all samples were combined to obtain the expression matrix. Final expression matrix was calculated by cross-species gene expression analysis as reported previously[12]. The expression values from mouse, rat and pig were transformed separately into relative abundance values: for each gene, the relative abundance value is the expression value divided by the mean of expression values within the same gene across samples in the same species. The final expression matrix was subjected to hierarchical clustering using R software. Development stage (PE, PrE, TE, VE and EPI)-specific genes were selected to do the subsequent analyses. They were mapped to the final expression matrix to do the PCA and heatmap analysis with R software.

**Generating of a GFP-KI reporter.**

In order to establish green fluorescent protein (GFP) gene-based reporter XEN cell line, we used a site-specific knock in (KI) Ossabaw fetal fibroblasts. In order to facilitate KI at high frequencies, we have used a combination of small molecule inhibitor of NHEJ pathway (SCR7)[13] and a pre-complexed Cas9 protein and sgRNA RNP complex to KI a ubiquitous promoter (UBC) driven GFP (Sanger Institute) downstream of a ubiquitously expressed *COL1A1* locus to ensure stable expression of transgenes. After a day of transfection, the GFP-positive (GFP+) cells were sorted by flow cytometry (Becton Dickinson, Franklin Lakes, NJ, USA) and GFP+ single cells were replated into wells of a 96-well plate for expansion. After 10–15 days, individual colonies were washed, suspended in 20 μL of lysis buffer (50mM KCl, 1.5mM MgCl2, 10mM Tris pH 8.0, 0.5% NP-40, 0.5% Tween-20 and 100 μg/mL proteinase K) and incubated for 1 h at 65˚C followed by heating the mixture at 95˚C for 10 min to inactivate the enzymes. The cell lysates (2 μL) were directly used as a template for PCR with screening primers (sFig. 1). Using this approach, we have identified >60% of the clonal lines showing stable integration of the transgene. The targeted-clones (hereafter called pCOL1A:GFP) with a strong and consistent fluorescence intensity as determined by fluorescence microscopy were frozen in 92% FCS and 8% DMSO, prior to use as nuclear donor cells. Using GFP labeled XEN cells, live animals were generated by SCNT.

**Chimera assay**

For lineage tracing of injected XEN cells, a total of eight reporter XEN cell lines were established from cloned blastocysts (Day 7 to 8), using GFP KI fetal fibroblasts (pCOL1A-GFP #3 and #6). A candidate female pCOL1a-GFP XEN cell line (Xnt pCOL1A:GFP#3-2) with stable expression of GFP and XEN markers was used for chimera testing. The cells were pre-treated with Rho Kinase (ROCK) inhibitor Y-27632 (10 μM; StemCell Technologies) for 2 hr and dissociated with Accutase at 38.5 °C for 5 min followed by gentle pipetting. About 3-4 small clumps (10–15 cells) were injected per blastocyst (*Supplementary Appendix, Fig. S3b*). After 20~24 hr of culture, injected blastocysts (n=94) were surgically transferred into the upper part of each uterine horn through needle puncture in recipients at days 5–6 of the estrous cycle (D0=onset of estrus; n=3). On day 15 after embryo transfer, the surrogate animals were euthanized to recover XEN-chimeras (XeC; embryonic day 21). A total of 25 fetuses were obtained after transfer and assessed macroscopically for viability and GFP expression. Fetuses that showed strong GFP expression in yolk sac (XeC#3-4) were cut sagittally; one half was used for histological analysis, whereas the second for DNA extraction. For detecting chimera contribution, gDNA were extracted from three parts of embryos: a small pieces of tissue at the posterior region of the fetus, yolk sac, and allantochorionic membrane. Embryos that were malformed or noticeably delayed (i.e. spherical and ovoid) were used only for gDNA isolation. The gDNA samples were subjected to PCR for chimera detection with genotyping primers (*Supplementary Appendix, Table S1*), and qPCR was performed for the detection of knock-in allele and chimerism rate. Prior to use in the qPCR analysis, the dynamic range of qPCR primers were validated (amplification efficiency >90%). The GFP labeled pXEN cell line (Xnt pCOL1A:GFP #3-2) was used as a positive control (GFP+, 100%) and a non-GFP XEN cell from parthenote embryo (Xpg#1) served as a negative (GFP-, 0%) control for investigating % chimerism. Relative expression was calculated using the comparative $2^{-\Delta\Delta Ct}$ method. qPCR was performed in triplicate. Cycling conditions for both GFP and reference (ACTB and YWHAZ gene) products were 10 min at 95°C, followed by 40 cycles of 95°C for 15 sec, and 60°C for 1 min. The primers used in qPCR are listed in *Supplementary Appendix, Table S1*.

**Teratoma Assay**

Immunedeficient-nude (BRG, BALB/c-Rag2$^{null}$ IL2rg$^{null}$; Taconic) and -scid (NIH-III, Cr:NIH-bgnu-Xid; National Cancer Institute) male mice were used to perform teratoma formation assay. Before transplanting, the pXEN cells were incubated for 2 hr in DMEM/F12 supplemented with Y27632 (10 μM). The cells were dissociated mechanically into small clumps, washed and suspended in 0.2 mL of mixture containing equal volumes of DMEM/F12 and Matrigel (Corning, MA, USA)[14]. With six pXEN cell lines, the cell suspensions (1 to 10 × $10^6$ cells) were subcutaneously injected into 6-8-week-old mice (*Supplementary Appendix, Table S2*). Mice were housed in specific pathogen–free conditions and were monitored for a minimum of 30 weeks.

**Appendix Table s1. Primers and Antibodies**

**qPCR for mRNA expression analysis**

| Gene | Forward | Reverse |
|---|---|---|
| *ACTB* | gtggacatcaggaaggacctcta | atgatcttgatcttcatggtgct |
| *AFP* | cacctttccaggttccagaa | aaggggtgccttcttgctat |
| *CK8* | tctgggatgcagaacatgag | ggctgtagttgaagcctgga |
| *CK18* | gcaagttctgtggacaatgc | gccagctccgtctcatactt |
| *CK19* | ctgaaggaagagctggccta | tcaacctccacactgacctg |
| *CXCR4* | cagcaagggtgtgagtttga | tccaaggaaagcgtagagga |
| *CDH1* | cacctcacgggaattgtctt | ttatcagcacccacgcaata |
| *DKK1* | aggctcttggaaccctgact | ccaaaggactcaaggcagag |
| *EPCAM* | ccaaaaggatggacctgaga | agcctgtagaccctgcattg |
| *FOXA2* | ataaggagggcaagggaaaa | agtcaaaattcgcaggtgct |
| *GATA4* | tctcggaaggcagagagtgt | caggcgttgcacaggtagt |
| *GATA6* | atcaccatcaccacccaagt | cgcgactctgtagactgtgc |
| *GPC1* | ccaggatgccagtgatgac | tggagcttttcttgctgacc |
| *GSC* | gaagccctggagaacctctt | cggtttttgaaccagacctc |
| *HNF4α* | ctcagcaacggacagatgtg | caggagcttgtagggctcag |
| *NANOG* | ccccttcttcaactcaaca | cttcaggcccataaacctca |
| *OCT4* | gctggagccgaaccccgagg | caccttcccaaagagaaccccaaa |
| *PDGFRα* | caggttggagggagatggac | agttgcggaggttggatt |
| *RN18S* | acaaatcgctccaccaactaaga | cggacacggacaggattgac |
| *SALL4* | caggagtaccagagccgaag | acctcgggagacttggactt |
| *SNAIL* | ttttcagcagccctatgacc | ccaggagagagtcccagatg |
| *SOX2* | aacagcccagaccgagttaa | gttgtgcatcttggggttct |
| *SOX7* | ggctagtgaaagccaactcg | tttgcctgccttgagagaat |
| *SOX17* | tggttgaatcttgaggtctgc | cagggtgtaggtgtgtgatga |
| *SPARC* | ggaccatcagtcctctggaa | agttctgcgtctcccaaaga |
| *TGFb1* | gtcttcttcggacgttaccg | gcatgaggaggaggaacaaa |
| *uPA* | aagggctctgacattccatg | ccggctcttacactgacaca |
| *VIMENTIN* | gtaccggagacaggtgcagt | ttccacggcaaagttctctt |

**qPCR for Chimerism analysis**

| Gene | Forward | Reverse |
|---|---|---|
| GFP | aagttcatctgcaccaccg | tccttgaagaagatggtgcg |
| YWHAZ | agtaggttgggctccttgacac | gccgactgtgactttaaggtgc |

**PCR for genotyping**

| Gene | Forward | Reverse |
|---|---|---|
| pCOL1A1 | gcatggagagaaggcatgat | |

| | | |
|---|---|---|
| Ubc promoter | | tcacagcgatccagaaagaa |
| NGN3 | caccagaccgagcagtcttt | ttggtgagtttcgcatcgt |

**Antibodies**

| Antigen | Antibody Source |
|---|---|
| AFP | Abcam (ab74663) |
| CDX2 | Abcam (ab76541) or Santa Cruz (sc19478) |
| EOMES | Santa Cruz (sc98555) |
| HNF4α | R&D Systems (AB41898) |
| SOX2 | Abcam (ab79351) or Santa Cruz (sc17320) |
| GATA4 | Santa Cruz (sc1237) |
| GATA6 | Santa Cruz (sc9055) |
| SALL4 | Santa Cruz (sc46045) |
| CDH1 | Antibodies-online (ABIN3209718) |
| GFP | Santa Cruz Biotech (sc9996) |
| LAMININ | Sigma Aldrich (L9393) |
| SOX17 | R&D Systems (AF1924) |
| SOX7 | R&D Systems (AF2766) |
| NANOG | Santa Cruz (sc33760) or Peprotech (500-P236) |
| OCT-4 | Santa Cruz (sc5279) |
| PCNA | Santa Cruz (sc56) |
| H3K27me3 | Epigentek (A4039-025) |
| VIMENTIN | Santa Cruz (sc6260) |
| Cytokeratin 8/18/19 | Abcam (ab41825) |
| Donkey anti-mouse | Santa Cruz (sc-2099 g) or Abcam (ab96878) |
| Donkey anti-rabbit | Santa Cruz (sc-2090) or Abcam (ab96894) |
| Donkey anti-goat | Santa Cruz (sc-2783) or Abcam (ab96935) |

**Appendix Table s2. Teratoma assay for determining the potency of pig XEN cells**

| pXEN lines | Injected cell No. | Stains | No. animals transplanted | No. teratoma developed |
|---|---|---|---|---|
| Xnt $_{Col1A:GFP}$#3-2 | $5 \times 10^6$ | BRG | 2 | 0 |
| | | NIH-III | 2 | 0 |
| | $10 \times 10^6$ | NIH-III | 1 | 0 |
| Xnt $_{Col1A:GFP}$#6-1 | $5 \times 10^6$ | BRG | 1 | 0 |
| | | NIH-III | 1 | 0 |
| Xvv#2 | $1 \times 10^6$ | BRG | 2 | 0 |
| | | NIH-III | 1 | 0 |
| | $10 \times 10^6$ | BRG | 1 | 0 |
| Xvv#9 | 1 X$10^6$ | BRG | 2 | 0 |
| | | NIH-III | 2 | 0 |
| | $10 \times 10^6$ | BRG | 2 | 0 |
| Xpg#1 | 1 X$10^6$ | BRG | 2 | 0 |
| | | NIH-III | 2 | 0 |
| | $10 \times 10^6$ | BRG | 2 | 0 |
| | | NIH-III | 2 | 0 |
| Xpg#4 | $5 \times 10^6$ | BRG | 1 | 0 |
| | | | 26 | 0 |

Six to eight week-old (BRG, BALB/c-Rag2$^{null}$ IL2rg$^{null}$ and NIH-III, Cr:NIH-bg-nu-Xid) male mice were used to perform teratoma assay. Before transplanting, cells were incubated for 2 hr in medium supplemented with Y27632 (10 μM) and were suspended with Matrigel matrix. Six XEN cell lines at passage 5-25 that were transplanted subcutaneously into 6-8 weeks old immunodeficient mice (n=26). Animals were monitored for 30 weeks or longer. However, teratoma formation in all six lines tested was not detected.

**Appendix Table s3. Integration of pXEN cells to pig blastocysts**

| Single / clumps | Stages | No. Injected | No (%). Blastocyst* | No. (%) contributed | into ICM | into TE |
|---|---|---|---|---|---|---|
| Single | Morula | 26 | 24 (92.3) | 9 (37.5) | 5 (20.8) | 4 (16.7) |
| | Blastocyst | 22 | 19 (86.4) | 9 (47.4) | 5 (20.8) | 4 (21.1) |
| Clump | Morula | 24 | 22 (91.7) | 17 (77.3) | 8 (36.4) | 8 (36.4) |
| | Blastocyst | 27 | 21 (77.8) | 18 (85.7) | 14 (66.7) | 4 (19.0) |

* The number in blastocyst injection is the number of re-expanded blastocysts on 2 day following injection. pXEN cells ($Xnt_{Col1A:GFP}$#3-2) were injected as individual cells or as small clumps after Accutase-treatment.

**Appendix Table s4.** A detailed analysis of distribution GFP+ cells in D21 pig chimeras

| Chimera No. | Yolk sac | | | | Amnion | Allantochorion | | | Embryonic | | |
|---|---|---|---|---|---|---|---|---|---|---|---|
| | Visceral endoderm | Mesothelium | Endothelium | Hematopoietic | | Mesenchyme | Epithelium | Hematopoietic | Ectoderm | Mesoderm | Endoderm |
| XeC#1-1 | - | - | - | - | - | - | - | - | - | - | - |
| XeC#1-2 | ++ | - | - | + | + | + | + | + | - | - | + |
| XeC#2-1 | - | - | - | - | - | - | - | - | - | - | - |
| XeC#2-2 | - | - | - | - | - | - | - | - | - | - | - |
| XeC#2-3 | + | - | - | - | ± | - | - | + | - | - | + |
| XeC#2-4 | ++ | - | - | + | - | + | - | ± | - | - | + |
| XeC#2-5 | + | - | - | ± | - | + | + | ± | - | - | - |
| XeC#2-6 | - | - | - | - | - | ± | - | - | - | - | + |
| XeC#2-7 | + | - | - | + | - | - | + | + | - | - | - |
| wt #4-2 | - | - | - | - | - | - | - | - | - | - | - |

Degree of GFP+ cells in chimeric conceptuses were expressed by - / ± / + / ++ means not/ weak/ moderate/ strong contribution, respectively. As a negative control wild type Day 21 embryo (wt#4-2) was collected from an artificially inseminated sow.

**Appendix Table s5.** ***In vitro*** **development of pig cloned embryos**

| Donor cells | Cell type | No. reconstructed | No (%). 2-4 cells | No (%). blastocysts |
|---|---|---|---|---|
| $FF_{wt}$#6 | Fibroblast | 25 | 22 (88.0) | 8 (32.0) |
| $FF_{Col1A:Attp}$#6-1 | Fibroblast | 107 | 87 (81.3) | 37 (34.6) |
| $FF_{Col1A:GFP}$#3 | Fibroblast | 40 | 34 (85.0) | 15 (37.5) |
| $Xnt_{Col1A:GFP}$#3-2 | XEN | 95 | 79 (83.2) | 36 (37.9) |
| Xvv#9 | XEN | 32 | 23 (71.9) | 14 (43.8) |

All the cells as nuclear donors were from the same origin ($FF_{wt}$ #6; a female Ossabow fetal fibroblast), except for Xvv#9 that was derived from an *in vivo* embryo (crossbred). There was no statistically significance between the groups.

### Appendix Figure legends

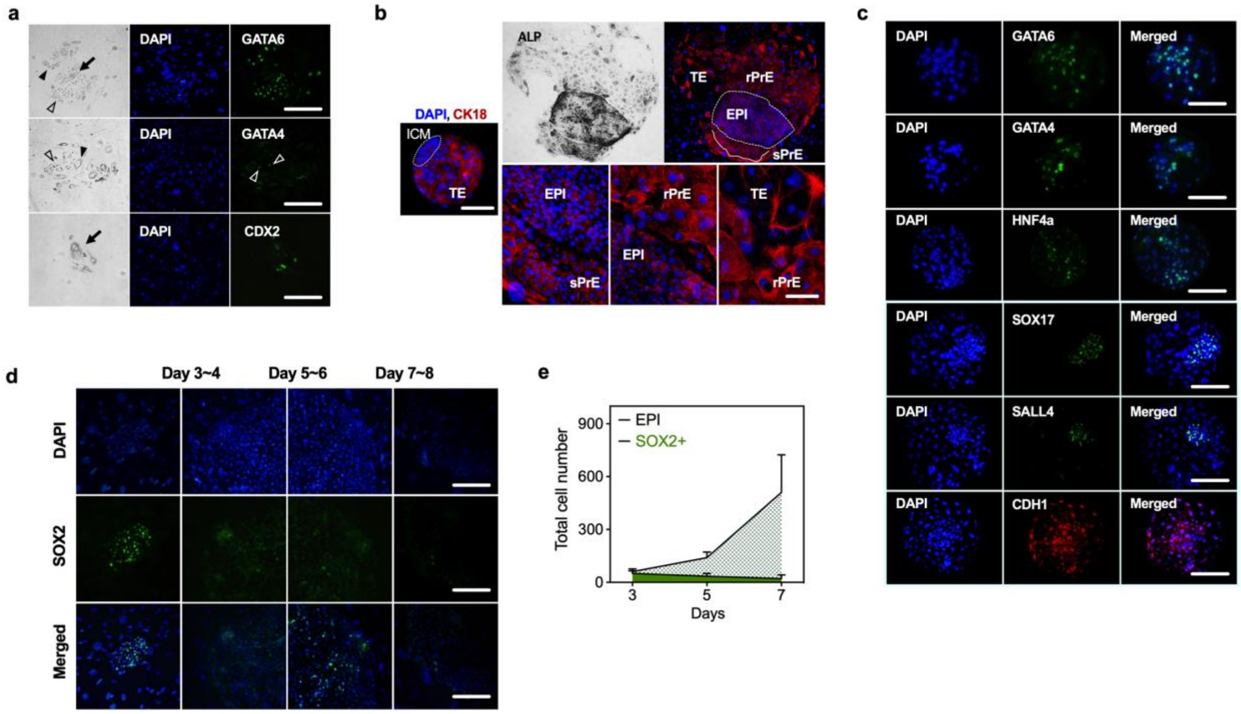


**Appendix Figure s1. Distinct subpopulations arise from the blastocysts outgrowth.**
(a) Phase contrast images and immunostaining of the primary outgrowth. In the primary outgrowth, GATA-positive large (filled arrowhead) and small (open arrowhead) round cells, and CDX2-positive trophoblast cells (filled arrow) were observed.
(b) Representative fluorescence images of CK18 in the blastocyst (ICM in dotted circle) and the primary outgrowth showing mixed populations, including large (rPrE) and small (sPrE) round cells.
(c) Representative fluorescence images of selected PrE markers in *in vitro* Day 7 blastocysts
(d) Representative immunostaining and (e) quantitation of the number of SOX2- positive nuclei in primary outgrowths cultured for 7 days.

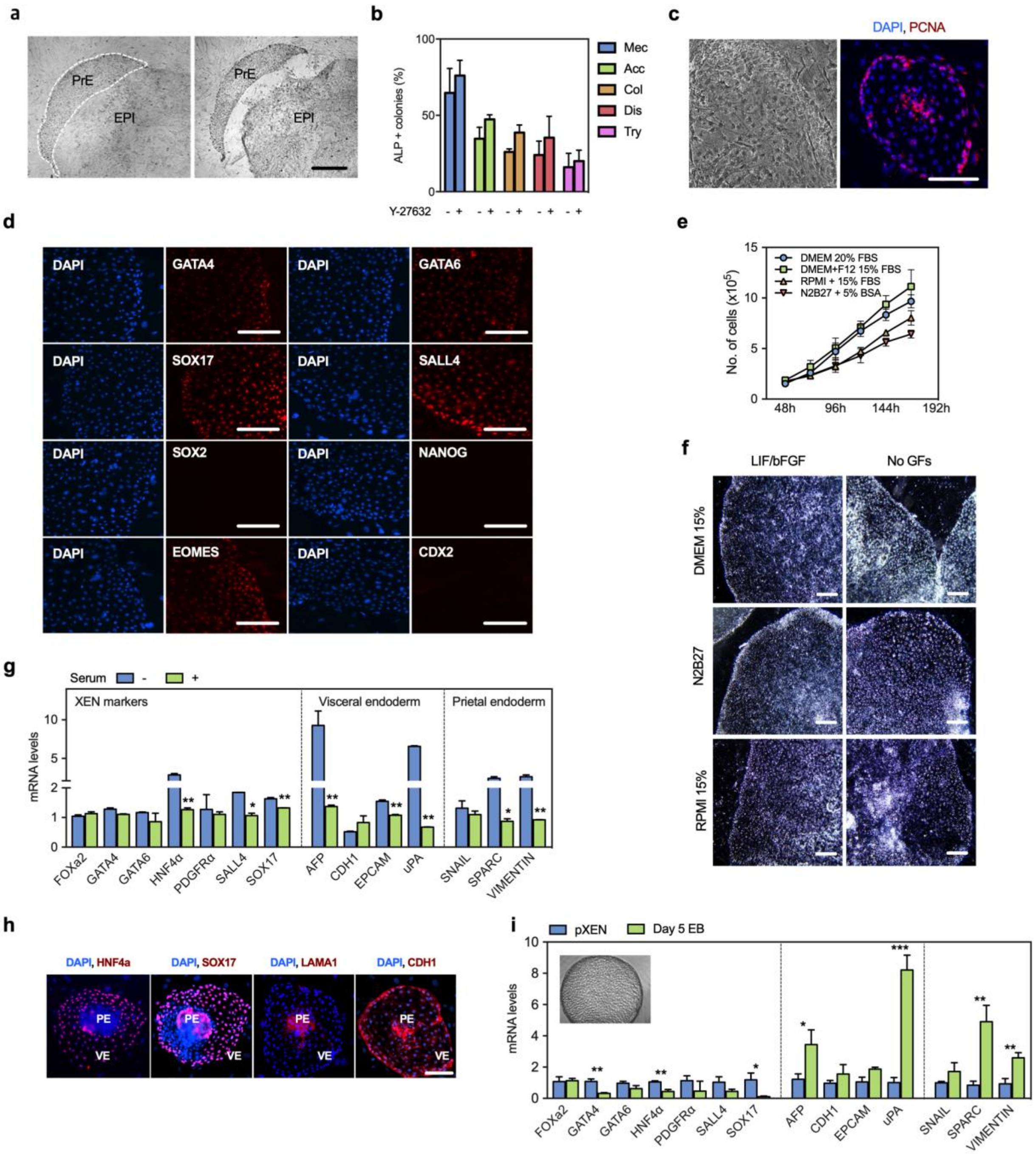


**Appendix Figure s2. Self-renewal of extra-embryonic endoderm XEN cells.**
(a) Representative bright-field images showing separation of PrE cells from the primary outgrowth after 7-9 days of culture.
(b) Efficiency of colony formation of pXEN cells passaged as clumps by mechanical (Mec) or enzymatic dissociation with Accutase (Acc), Collagenase IV (Col), Dispase (Dis), and Trypsin 0.5% (Try) in the presence or absence of ROCK inhibitor (Y-27632)
(c) Representative images of pXEN cells show the expression of proliferation marker, PCNA (right).
(d) Expression of the indicated markers in pXEN at passages 30-35.

(e) Effect of culture medium during propagation of pXENs. The cells were seeded in 6-well-plates at a density of $5 \times 10^4$ cells per $cm^2$ and estimated 48h after a lag period following passage. Data are presented as means ± s.d. (n = 3).
(f) Representative bright-field images of pXEN in different culture mediums.
(g) qPCR analyses with total RNA isolated from pXEN cells grown in either the presence or absence of LIF/bFGF for 4 days. ACTB was used as a loading control. The values are represented as mean ± s.d. (n = 3).
(h) Expression of the indicated markers in pXEN cells. Two subpopulation frequently observed during propagation of pXENs.
(i) qPCR analyses with 5-day-old EBs derived from pXENs using the hanging drop method. ACTB was used as a loading control. The values are represented as mean ± s.d. (n = 3).

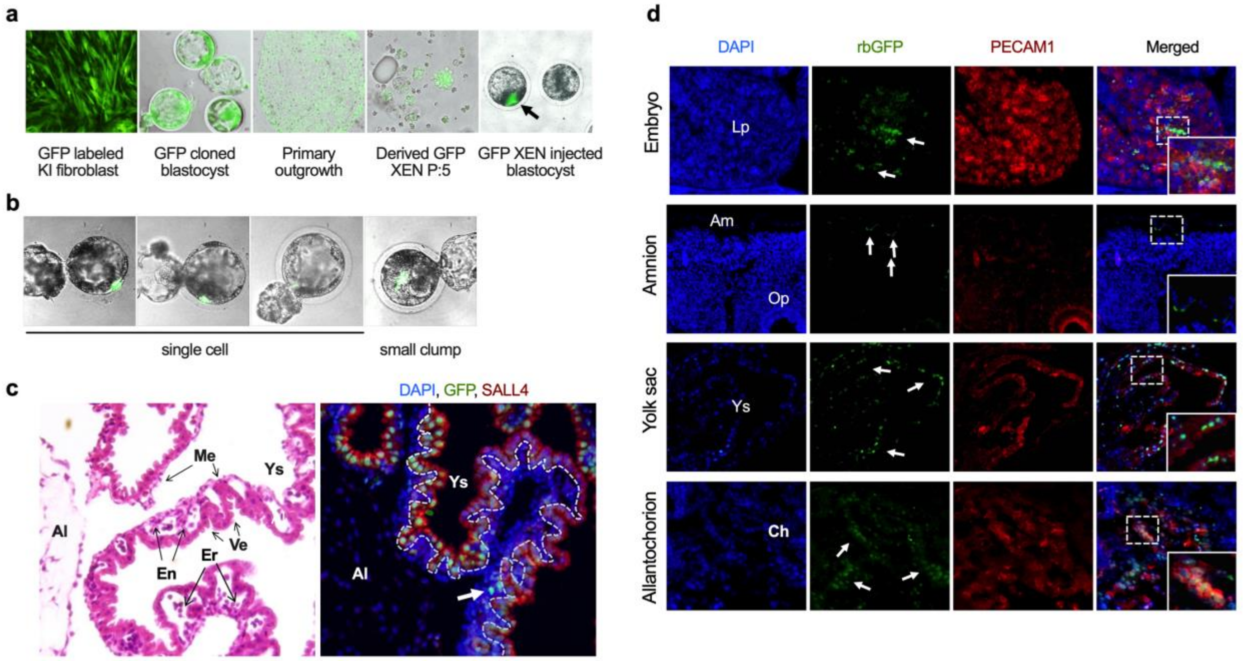


**Appendix Figure s3. Chimeric contributions of pXEN cells in embryo**
(a) Representative images of generation of GFP-labeled XEN (filled arrow) cell line and chimeric embryos (b) Representative sagittal sections showing hematoxylin and eosin (H&E) stains and immunofluorescence for GFP (green) and SALL4 (red) in a chimeric D21 yolk sac. In the left panel, the yolk sac consists of 3 thin cellular; the inner surface of the mesothelium (Me) the outer layer of visceral endoderm (Ve), the yolk sac cavity with primitive erythrocytes (Er) surrounded by a layer of endothelial cells. In the right panel, section was immunostained with anti-GFP (green) and anti-SALL4 antibodies, which were present in the visceral endodermal layers (dotted line). A few GPF-positive cells were observed in the primitive erythrocytes (arrow).
(c) Section was immunostained with anti-GFP (green; arrow) and anti-PECAM1 antibodies showing that cells from GFP-pXEN contribute to embryonic tissues and fetal membranes in a D21 chimera (#1-2). The area in the dashed box are displayed at a higher magnification. Nuclei were stained with DAPI (blue). Ch, chorion; Lp, liver primordium; Ys, yolk sac; Am, amnion, Op otic pit.
